# A simple correction to adjust for sampling biases in phylogeographic discrete trait analysis

**DOI:** 10.1101/2023.11.21.568020

**Authors:** David Jorgensen, Margarita Pons-Salort, Nicholas C Grassly

## Abstract

Discrete trait analysis (DTA) methods are widely used to infer patterns of movement between discrete geographic locations from genetic sequence data as they are simple and fast to implement. With the increasing interest in phylogeography for pathogen genomic epidemiology, accounting for and understanding the limitations of the methods used is becoming increasingly important. Applying DTA to location makes several strong simplifying assumptions, including the assumption of equal sampling probability across locations. This may lead to biased estimates of migration rates and ancestral locations where sampling rates vary. Here we investigate these biases using simulated migration and transmission data. We propose a simple correction to DTA for unequal sampling probability across locations when sampling rates are known or can be estimated. This correction to the migration rate matrix can be applied only to phylogenetic branches leading to sampled tips or to the whole phylogeny. We assess the performance of this correction using simulated data for a range of endemic transmission scenarios with varied sampling and migration rates. In a simple, two-location, case the correction performs significantly better than a standard DTA method in three of the four sampling rate distributions investigated. Applying the correction across only the tips of the phylogeny and over the full depth of the tree gives similar results, with the correction only at the tips performing marginally better in this simple two-location simulation. As migration is a key driver of infectious disease spread, improving estimates of migration with simple phylogenetic methods is a valuable development in infectious disease phylogeography.

## 1 Introduction

With the increasing availability of high quality and geographically diverse genetic sequence data from emerging and endemic diseases, phylogeographic analysis is becoming increasingly popular to estimate likely sources and routes of viral transmission. Recent studies have applied phylogeographic methods to study Influenza, Rabies, HIV, Ebola, Zika and the SARS-CoV-2 pandemic, amongst other viral infections [1, 2, 3, 4, 5, 6, 7, 8, 9]. Observed viral sequences are often annotated with location information and phylogeographic research aims to combine this information with the inheritance information provided by the genetic sequences themselves to infer the likely location of unobserved ancestral virus and the rate of movement of virus between locations. A number of methods have been developed to reconstruct ancestral states on a fixed phylogeny [10, 11]. Traditionally, this mapping was carried out using parsimony methods, minimising the number of state transitions required to explain the observed traits at the tips of the phylogeny [11]. This can be misleading where rates of migration are high or state exchange probabilities are unequal, both important considerations in infectious disease phylogeography [12, 13].

A simple and widely used alternative is discrete trait analysis (DTA) (also often referred to as ‘mugration’ as the migration process is modelled using similar methods to those used to model mutation, with each location modelled as a state of the discrete trait) [13]. Using a probabilistic method allows a model of location or ‘state’ change to be explicitly specified and transition rate parameters to be estimated. DTA does not directly use the location data during inference of the phylogeny, but infers migration along the branch lengths and estimates the geographic location of internal nodes. This allows the evolution of the discrete trait to be informed by both a model of dispersion patterns over time and the structure of the phylogeny previously inferred from the genetic sequence data [13]. DTA methods typically employ continuous-time Markov chain models from the ‘Mk’ family of Markov models of discrete character evolution [14]. Pagel (1994) was the first to propose the use of an Mk model for discrete trait data with a probability distribution of character states inferred at each internal node on the phylogeny, the basic structure used for contemporary maximum likelihood discrete trait analysis [15].

Most discrete trait models consider ‘reversible’ transition rates, where the forward and backward rate of movement between a pair of discrete states are equal, as these are more commonplace in the substitution models already widely implemented in phylogenetic software. The TreeTime mugration method of Sagulenko et al., for example, implements the generalised time reversible model of Tavaré to estimate migration and forms the basis of the internal node location estimates in the popular, open-source, Nextstrain pipeline [16, 17, 9]. Here, we consider a non-reversible or ‘all-rates-different’ (ARD) Mk model applied to a single discrete trait, location. The aim of employing a non-reversible model is to better capture asymmetric migration patterns. This has an associated cost of doubling the number of transition rate parameters which need to be inferred over a reversible model.

When modelling human movement in epidemiology it is common to consider either the radiation or gravity models, where movement of individuals is incorporated as a directional process [18]. These models were developed to account for short-term movement of individuals such as commuting and retail, key drivers of the spread of infectious diseases between populations. This directionality is also an intuitive property of medium-term human migration with certain regions gaining and losing population over time within a country [19]. This differs from traditional molecular ecology where we are considering the development of traits over ‘ecological time’, measured in generations, and drivers of individual movement are of less importance to trait reconstruction [20]. As the spread of human viruses between populations is driven primarily by the movement of individuals between locations, it follows that movement of virus between populations will be similarly directional. Directional discrete trait models are implemented in the ace function from the R phylogenetics package ape and are supported for discrete trait reconstruction in the Bayesian phylogenetic software BEAST [21, 22, 23].

Although DTA is computationally efficient and easy to implement with a simple maximum likelihood implementation of Felsenstein’s pruning algorithm, the method was designed to model the substitution of alleles at a genetic locus. Here we instead aim to discern the movement of individuals between discrete locations [10]. The primary difference between the two is that phenotypes used in traditional discrete trait models are directly linked to the underlying genetic data informing the phylogeny whereas genetic data has no direct causative effect on the location of a sample.

Assuming that migration is analogous to mutation inherits assumptions from mutation models at odds with traditional models of migration, outlined succinctly by De Maio et al. [24]. Briefly, DTA assumes i) the shape of the phylogeny is independent of migration, ii) sampling intensity in each state is proportional to the infected population in that state at the time of sampling, iii) prevalence of states can drift with states able to become fixed or extinct over time, iv) any states not sampled at the tips are not able to be inferred as ancestral locations. These factors can lead, amongst other issues, to biased estimates of migration rates and the location inferred at internal nodes where there is unequal sampling between states [25].

Alternative phylogeographic methods which are better suited to structured populations (such as the spatial and temporal subdivisions present in epidemiological models) are also becoming more widespread. One such model is the structured coalescent, introduced by Hudson and Notahara in 1990 and based on the earlier coalescent method of Kingman, 1982 which is implemented in a range of phylogenetic software [26, 27, 28, 29]. The structured coalescent is more appropriate than discrete trait analysis for structured populations as it incorporates fixed effective population sizes, allowing sampling rates to differ across populations, and allows the population structure to affect the shape of the inferred phylogeny. The structured coalescent can still result in improper posterior densities for migration rate and population size parameters where sensible bounds are not given or uninformative priors are used for demographic parameters [30, 31]. Full structured coalescent models can also be slow to, or fail to, converge where more than three or four discrete states are used, which can also be an issue when incorporating complex population structure [31, 32]. For this reason, and to incorporate predictor variables, a number of approximations and extensions to the structured coalescent have been proposed in recent years. These include the approximations proposed by Volz, De Maio et al. and Muller et al. which avoid the sampling of migration histories in the structured coalescent by integrating over possible migration histories [33, 24, 32]. Differences between these approximations are described in Muller et al. 2017 [34]. Despite these simplifying assumptions, there are still limitations to the numbers of sequences and locations which can be considered with these methods, in the hundreds of sequences and fewer than 10 discrete location states. This precludes their application to large datasets and complex populations. Both the DTA and the marginal approximation of the structured coalescent proposed by Muller et al. have also been extended to allow additional migration predictor information to be provided with a generalised linear model [35, 32]. This can improve inference estimates where demographic or distance parameters, for example, affect the movement probability between locations.

Other methods such as the birth-death process and the generalised coalescent can be used to combine epidemiological models of infection and recovery with genetic data and analyse phylogenies constructed with data from structured populations [36, 33]. These both allow more complex models of birth, death, migration and sampling to be specified, although these models may also be biased and can be slow and difficult to implement. Alternatively, location data can be treated as continuous rather than discrete information, which has the benefit of not requiring homogeneous discrete regions to be defined [37, 38, 39]. Defining these regions is a key step in any discrete phylogenetic approach, with results strongly dependent on the regional boundaries and extent of the study area considered [40, 41]. As DTA continues to be used widely due to the speed and scalability of the method to large numbers of states, context-specific amendments are important to produce reliable results. Here we propose a simple change to the transition matrices central to the likelihood calculations used in discrete trait analysis to correct for unequal sampling rates between locations.

## 2 Methods

Simple maximum likelihood discrete trait analysis uses k-state Markov models (Mk models) of migration between locations [14]. Using a Markov transition matrix and Felsenstein’s tree pruning algorithm it is possible to estimate the likely locations of internal nodes in a phylogeny. This process treats migration akin to mutation of a discrete trait, hence the name ‘mugration’ is often used to describe discrete trait analysis applied to migration [10, 42, 24]. These Mk models are traditionally used in molecular ecology as genetic substitution models, with the simplest equal rates and equal frequency model introduced by Jukes and Cantor in 1969 [43]. Here we consider the ARD Mk model where the migration rate from location i to j can differ from the reverse rate from j to i. Models of this type are not typically used as substitution models, although they were introduced for this use by Barry and Hartigan in 1987 [44, 45]. Statistical identifiability issues for tree generation with the non-reversible general Markov model were described by Zou et al. in 2011 [46]. In the case of DTA, as we are considering changes in a discrete trait independently of the process of tree generation, the non-reversible model can be used and is offered as a method in popular phylogenetic software [21].

### 2.1 General methodology

In an Mk model, the instantaneous rate of change between state *i* and *j* is denoted by *q*_*ij*_. General Markov models of discrete character evolution with *k* states can be represented with a transition rate matrix, Q [14, 47] (Eq. 1):

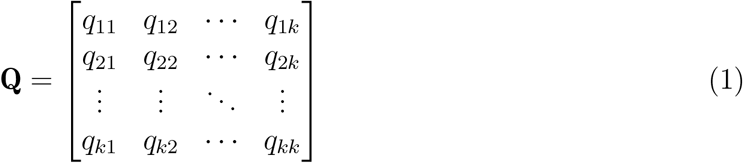

Where each element on the main diagonal is equal to the negative of the sum of all other elements on the same matrix row (*q*_*kk*_ = − Σ_*i ≠k*_ *q*_*ki*_). The transition rate matrix can be used to compute the transition probability matrix P(*t*) for a time interval t by taking the matrix exponential [48](Eq. 2). The resulting matrix (P(*t*)) is a right stochastic Markovian transition matrix.

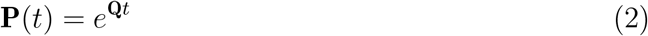

The likelihood for an ancestral node is given by the product of this transition matrix (P(*t*)) and a vector (*v*_*desc*_) corresponding to the location probability of each descendent node. In the case of a tip, where the sampling location is known, this probability is 1 for the sampled location and 0 for all other locations. As each ancestor has two descendants, we need to calculate P(*t*)*· v*_*desc*_ for each descendent branch. The product across these two branches gives the overall likelihood at the ancestral node. Once the ancestral node likelihood is assigned the descendent branches from that node can then be ‘pruned’ and the likelihood calculated recursively up the tree using Felsenstein’s pruning algorithm [10]. The overall tree likelihood is calculated by taking the product of these values across all internal nodes.

### 2.2 Proposed corrections

We propose and test the performance of four corrections to attempt to account for sampling rate differences between locations in DTA. When testing the corrections, we consider sampling rate as the proportion of all currently infected individuals sampled per unit time at endemic equilibrium. The four corrections use a similar method to the equilibrium base frequency vector supplied in the generalised time reversible DNA evolution model of Tavaré [16]. These four proposed corrections are made up of two related corrections applied only at branches leading to the tips of the phylogeny (Fig. 1) or over the full depth of the phylogeny, as in the GTR model of Tavaré.

**Figure 1:**
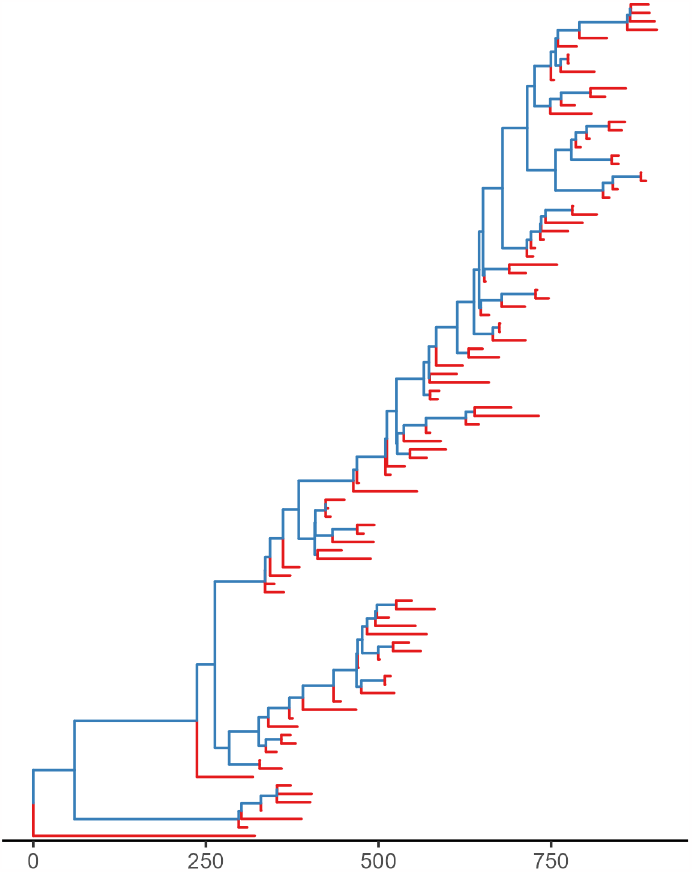
Example of a 100-tip phylogenetic tree with the terminal branches coloured red to show the branches over which the sampling rate correction is applied in the ‘tip-only’ corrections. Internal branches where the Q matrix is left unaltered are coloured blue. The x-axis shows the number of days from the time of the common ancestor (at zero) of the simulated samples (the root of the phylogeny).

The first correction applies the sampling rate directly, with a diagonal matrix of sampling rates multiplied to the Q matrix via matrix multiplication (Eq. 3).

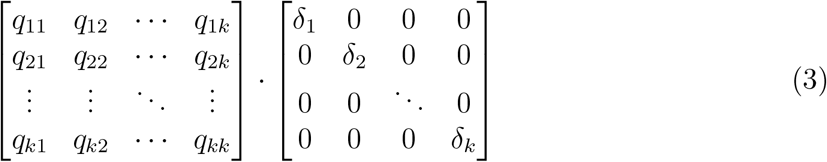

where each of the k rates of moving into a location i (*q*_*ki*_) are multiplied by the corresponding sampling rate in that location (*δ*_*i*_). We then set the diagonal elements to the negative of the sum of all other elements on the same matrix row (Eq. 4).

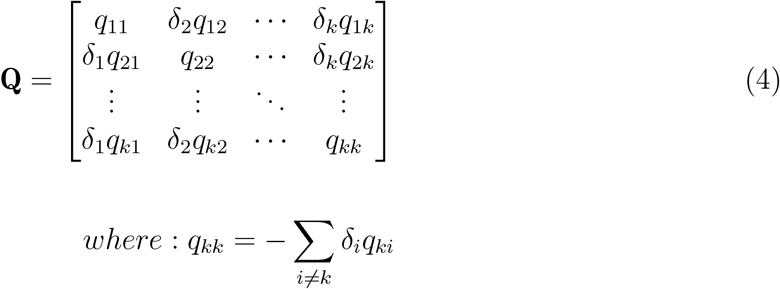

The second correction takes the additional step of normalising the sampling rates. We normalise the sampling rates to sum to one, akin to the treatment of equilibrium base frequencies in the GTR model. As the magnitude of the sampling rates is already reflected in the tree structure; with branch lengths and internal node positions in time influenced by the number of samples taken per day, this normalisation is designed to reduce the order of magnitude differences in sampling rates between scenarios to a set of relative rates on the same scale across scenarios. As all sampling rates are positive, the normalisation is carried out by dividing the rate in each location by the sum of rates across the k locations (Eq. 5) to give normalised sampling rates *ρ*_1_ *· · · ρ*_*k*_.

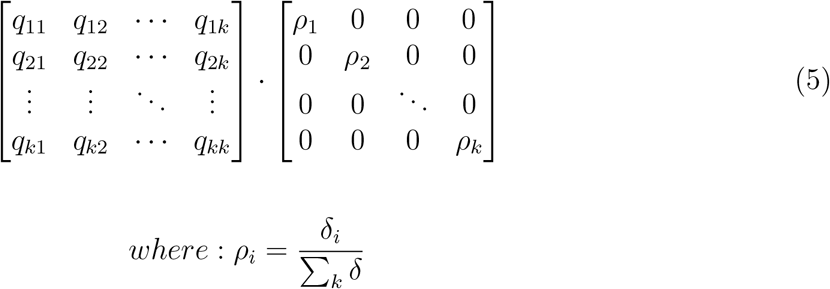

The diagonal elements of the Q matrix are given by the negative of the sum of all other elements on the same matrix row (Eq. 6), as in the previous proposed correction.

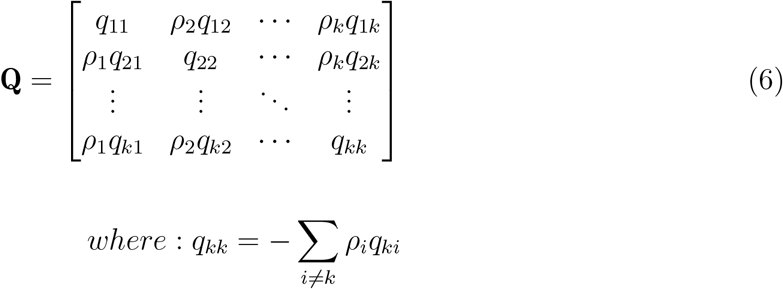

These two correction methods are applied either at branches leading to tips, as illustrated in Fig. 1, or over the full depth of the tree, giving the four proposed corrections. We propose these two alternative implementations as, where migration and mutation occur at a higher rate than sampling, it holds that the phylogeny produced will have a more diverse sample at the tips than where sampling rates are higher than migration and mutation rates. We posit that where a more diverse sample is collected, and thus the root of the tree is deeper and internal branches are longer, sampling rate differences at the tips will have a greater effect on the inferred location of the nodes directly ancestral to the tips than the deeper tree. Viceversa where the sample is less diverse, and the root of the tree is less deep in time we suggest that sampling may have a greater effect on the structure of the full phylogeny. Where the correction is applied only at branches leading to tips the general methodology is followed for internal branches, with the Q matrix applied as given in equatxion 1. Work by Faria et al. applied a similar branch partitioning approach to account for differences between successful host-shifts and dead-end cross-species transmissions in a model of zoonotic infection[49].

Applying the adjustments prior to calculating the diagonal elements of the Q matrix preserves the properties of the transition rate matrix as outlined in Norris 1997 [50] (Eq. 7), namely:

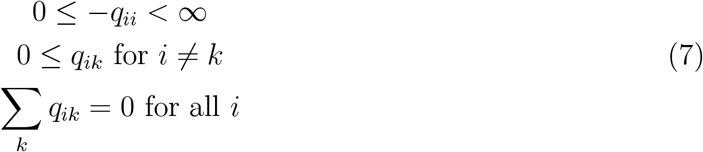

Giving rows which sum to zero and no negative off-diagonal or positive diagonal elements.

The proposed corrections are not perfect as sampling events also influence the tree structure and the timing of internal nodes. In the DTA, phylogenies are estimated via a separate process and the topology of the phylogeny is not updated to account for the inferred migration model. This means that the influence of migration on tree topology cannot be accounted for, and this would require a more complex phylogeographic model.

### 2.3 Tree simulation

To test the proposed model corrections, we consider a range of simulated phylogenies with different migration and sampling rates. The simulated phylogenies are produced under a multitype birth-death (MBD) model using the BEAST2 package MASTER [51, 52].

A simple epidemic model is simulated with a single infection seeded in one of the locations at time zero. For simplicity we assume that migration occurs at a constant rate over time and that there is no substructure within the states, with all individuals within each state having the same fitness and same probability of being sampled. The model is allowed to progress for 10000 time-steps (days) without sampling to ensure an endemic equilibrium is reached prior to sampling. A sampling compartment is used with no outflows to ensure sampled individuals are an endpoint for transmission. This is a simplification of real sampling dynamics for mathematical convenience, preventing a sampled lineage from being directly ancestral to another sample which would result in a zero length branch. When reconstructing the tree, the time of movement into the sampling compartment is taken as the time of the sampled tip. Recovered individuals are considered immune and no longer contribute to the transmission as infectors or susceptible individuals. Each sampled phylogeny is taken from an independent simulation and contains 100 sampled tips across the two sampled populations, regardless of sampling rates. We do not simulate genetic sequences for these phylogenies, instead we use only the simulated labelled trees and branch lengths. In other words, we do not account for phylogenetic uncertainty relating to the quality, length, and characteristics of any sequence.

Figure 2 shows a flowchart representation of one of the populations included in the MBD model. Rates shown in Figure 2 but not in table 1 are varied to test the effectiveness of the model corrections under different scenarios outlined below.

**Table 1:**
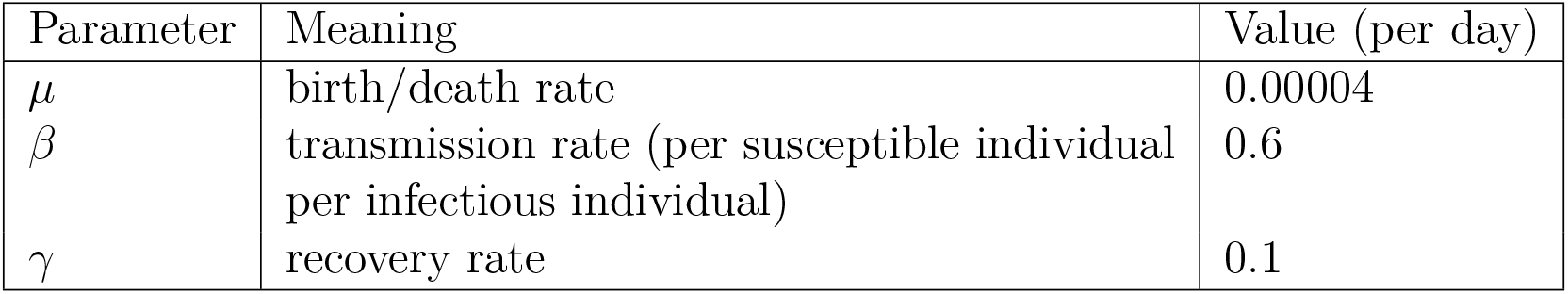
Input parameter values fixed across models. Beta and gamma are set to give an R0 of 6 and infectious period of 10 days and the birth/death rate is set in days to give a life expectancy of 68 years.

**Figure 2:**
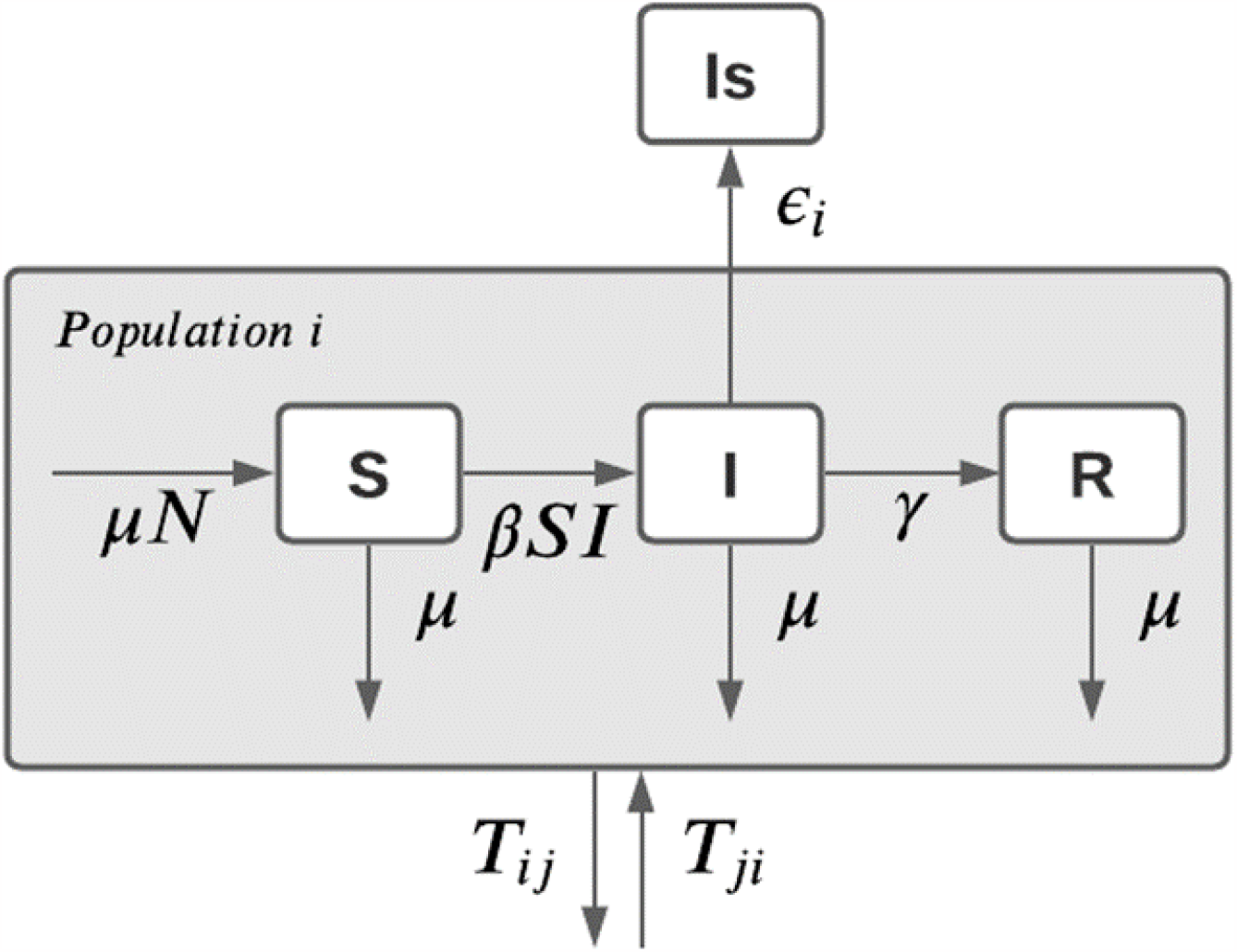
Structure of the simple SIR model implemented in MASTER in one of two populations. Additional populations are added as identical modules with additional migration rates to allow movement between all populations. The compartments represent Susceptible (S), Infected (I) and Recovered (R) individuals with the additional Is compartment representing sampling with removal.

### 2.4 Modelled scenarios

For ease of computation and understanding, we test two-state implementations of the correction methods. Further tests with more states will be required to further validate the correction methods for use with greater numbers of states, the primary use case for DTA. For each of the two-state MBD models we consider a range of relative migration and sampling rates between the two modelled locations. We test relative rates up to 4 times higher in each location over the other by randomly selecting migration rates between 0.005 and 0.05 per person per day to satisfy the relative rates we are aiming to reconstruct.

To test the methods under different sampling scenarios we vary the sampling and transmission rates. These scenarios are the “high sampling rate” scenario, where the sampling rates are an order of magnitude higher than migration rates; the “medium sampling rate”, where the migration and sampling rates are on the same order of magnitude; the “low sampling rate”, where the sampling rates are an order of magnitude lower than the migration rates; and the “transmission difference” scenario where sampling rates are the same as in the low sampling rate scenario but transmission in population 2 is twice as fast as in population 1, leading to a larger number of infected individuals at equilibrium in this population (Table 2). This covers a realistic range of relative migration and sampling rates between populations.

**Table 2:** Migration and sampling rate ranges for each of the modelled scenarios.

| Model scenario | Migration rate range | Sampling rate range |
| --- | --- | --- |
| High sampling rate | 0.005 to 0.05 | 0.05 to 0.5 |
| Medium sampling rate | 0.005 to 0.05 | 0.005 to 0.05 |
| Low sampling rate | 0.005 to 0.05 | 0.0005 to 0.005 |
| Transmission difference * | 0.005 to 0.05 | 0.0005 to 0.005 |

\* The transmission difference scenario has identical migration and sampling rates to the low sampling rate scenario but different transmission rates between locations (Transmission difference: $\beta_1 = 0.6$ and $\beta_2 = 1.2$ ; Low sampling rate: $\beta_1 = \beta_2 = 0.6$ )

### 2.5 Fitting the model

We test the adjusted model by adapting the ‘ace’ function from the R phylogenetics package ‘ape’ [21]. This method uses maximum likelihood optimisation which, although simple and fast to implement, can often have issues with the Mk model. Previous work has reported a flat or ridged likelihood surface for a two-state model, making it impossible to reconstruct exact values for the *q* parameters [24, 47]. As we find similar issues with the reconstruction of directional migration rates, we reparameterise the migration model to estimate a relative migration rate between a pair of locations rather than attempting to estimate each migration rate. In order to estimate a relative rate directly from the likelihood maximisation, we amend the Q matrix (Eq. 8):

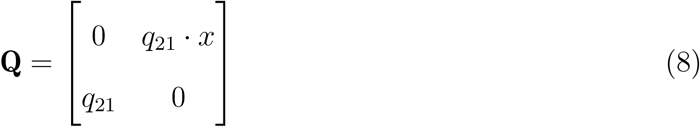

Giving a base rate of migration from location 2 to location 1 (*q*_21_ in eq. 8) and a rate relative to this base rate for the reverse migration (*x* in eq. 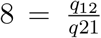). This minimises convergence issues resulting from collinearity between forwards and backwards rates as the relative rate remains consistent for a range of base rates along a ridge, meaning a constant relative rate should be returned as long as likelihood maximisation results in a point on the ridge. Although this change does not lead to a convex likelihood surface, the ideal scenario, it does allow for comparison between model fits (Fig. 3). This change is carried out prior to applying corrections and setting the diagonal elements, as outlined in equations 3 to 6.

**Figure 3:**
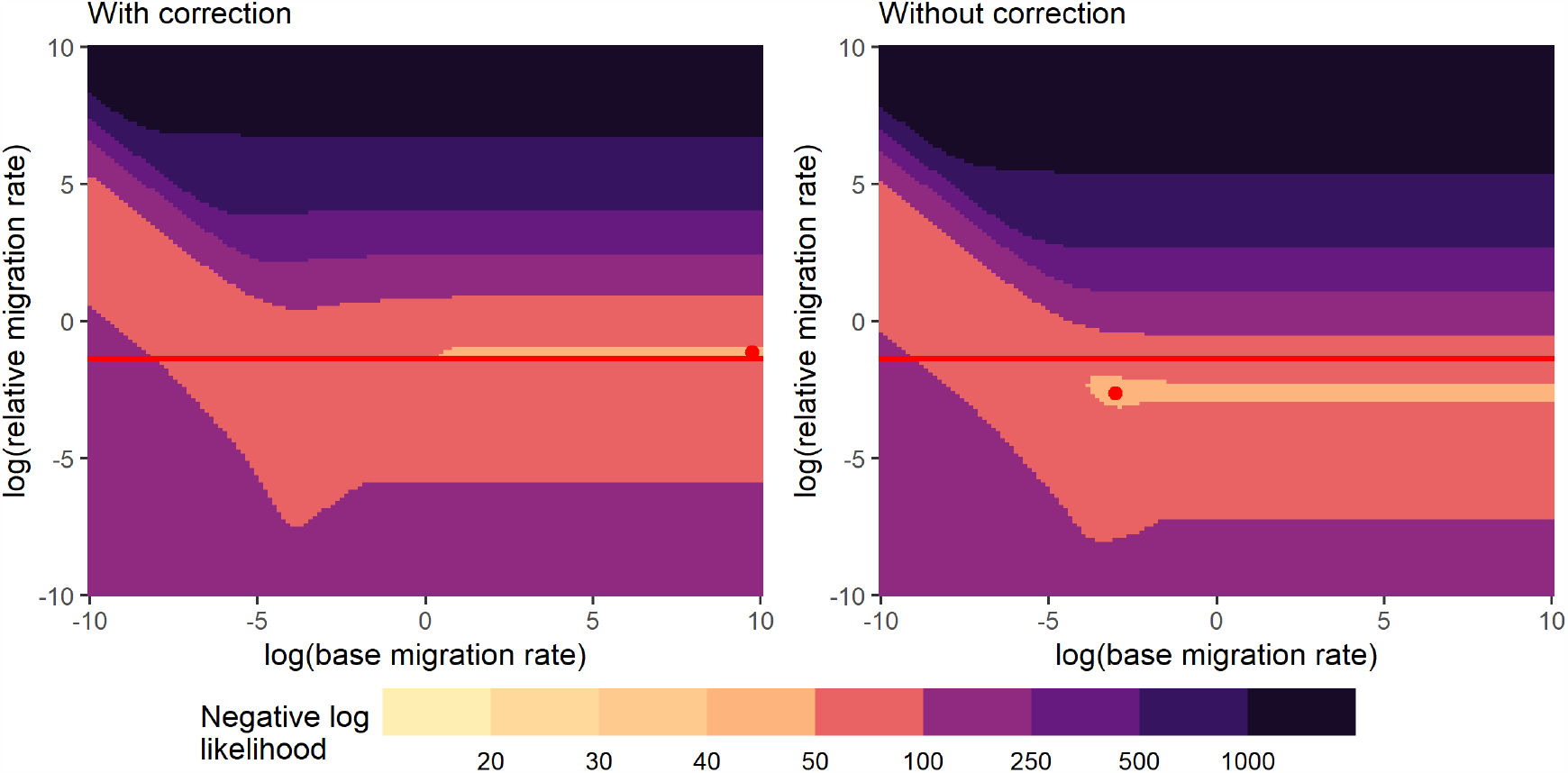
Likelihood surfaces for the base and relative migration rate. These are plotted on a log scale as the maximum likelihood estimation is performed on log-transformed parameters. This plot highlights the ridged likelihood surface and the improvement in maximum likelihood estimation with the proposed correction. This surface is for a single tree with simulation migration rates of 0.01 (pop 1 to 2) and 0.04 (pop 2 to 1) and sampling rates of 0.0039 (pop 1) and 0.000975 (pop 2). The surfaces are split into bins, with bounds shown in the figure legend. The lightest colour represents the maximum likelihood (minimum negative log likelihood) region. The horizontal line on each plot represents the true relative migration rate we are attempting to reconstruct (0.01/0.04 = 0.25, *log*(0.25) = − 1.39) and the red dot shows the location of the maximum likelihood estimate from the surface.

As relative rates have a lower bound of zero and, where both rates are equal, have a value of 1, the natural logarithm of the relative rate will approximately follow a normal distribution. We perform likelihood maximisation on the log-relative-rate, allowing standard errors to be estimated on the normal distribution, resulting in correctly distributed approximate confidence intervals when transformed back to a linear scale. This transformation has the added benefit of a *±*∞ scale, meaning a wider range of likelihood maximisation techniques can be used. We can reconstruct the approximate standard error for this log-relative-rate from the hessian of the maximum likelihood region of the likelihood surface. Here we use the Hessian method implemented in the ‘optim’ function from the ‘stats’ R package. Unfortunately, the eigenvalue-based solution used assumes a convex likelihood surface around the maximum likelihood point estimate and can fail to reconstruct a standard error where the likelihood surface is flat in one or more directions, as is the case with the models tested. Where confidence intervals could not be reconstructed with either the corrected or uncorrected methods, or both, the phylogeny was dropped for mean squared log error (MSLE) and confidence interval comparisons.

We assess the ability of the corrections to reconstruct relative migration rates by comparing the MSLE between the true and estimated relative migration rates across all simulations. Significance is assessed with a p-value threshold of 0.001 due to the very large sample size of 10,000 trees used.

## 3 Results

### 3.1 Correction selection

For the corrections using the exact rather than the normalised sampling rate, the overall MSLE was 0.26 applied across all nodes and 0.19 applied only at the branches leading to tips. Where the sampling rates were normalised to sum to one, the overall MSLE was 0.18 applied across all nodes and 0.14 applied only at branches leading to tips. As the correction with normalised sampling rates applied to branches leading to tips performed best across all sampling scenarios, the results from this method are presented in the main text below. The normalised correction applied to the full depth of the tree performed better than either of the non-normalised corrections.

### 3.2 Likelihood surfaces

The likelihood surface for the migration rate from 2 to 1 and the relative rate of return (*q*_21_ and *x* in eq. 8 respectively) showed a repeated trend of heavy ridges, where the maximum likelihood relative migration rate is constant across a large range of base migration rates. This ridged maximum likelihood region is brought more into line with the true relative migration rate with the proposed normalised tip-only correction. An example of the performance of the correction for a single phylogeny presented in figure 3. In both the corrected and uncorrected cases shown in figure 3 the maximum likelihood ridge is consistent for high values of the log-transformed base migration rate, becoming distorted at lower values. This trend is seen over the range of sampling and migration rates investigated, although the effect is more pronounced with more heavily biased rates.

### 3.3 Relative migration rate estimates

#### 3.3.1 Low sampling rate and transmission difference scenarios

As the low sampling rate and transmission difference scenarios both use sampling rates an order of magnitude lower than the migration rates, we present the results together. The scenarios differ as the transmission difference scenario incorporates a 2-fold higher transmission rate in one population than the other (*β*_1_ = 0.6, *β*_2_ = 1.2) to assess the ability of the method to correct for sampling rate differences where transmission is unequal across regions (Table 2). Fig. 4 shows the proportion of model runs where the true relative migration rate was within the 95% confidence interval of the estimated relative migration rate under the corrected and uncorrected models.

**Figure 4:**
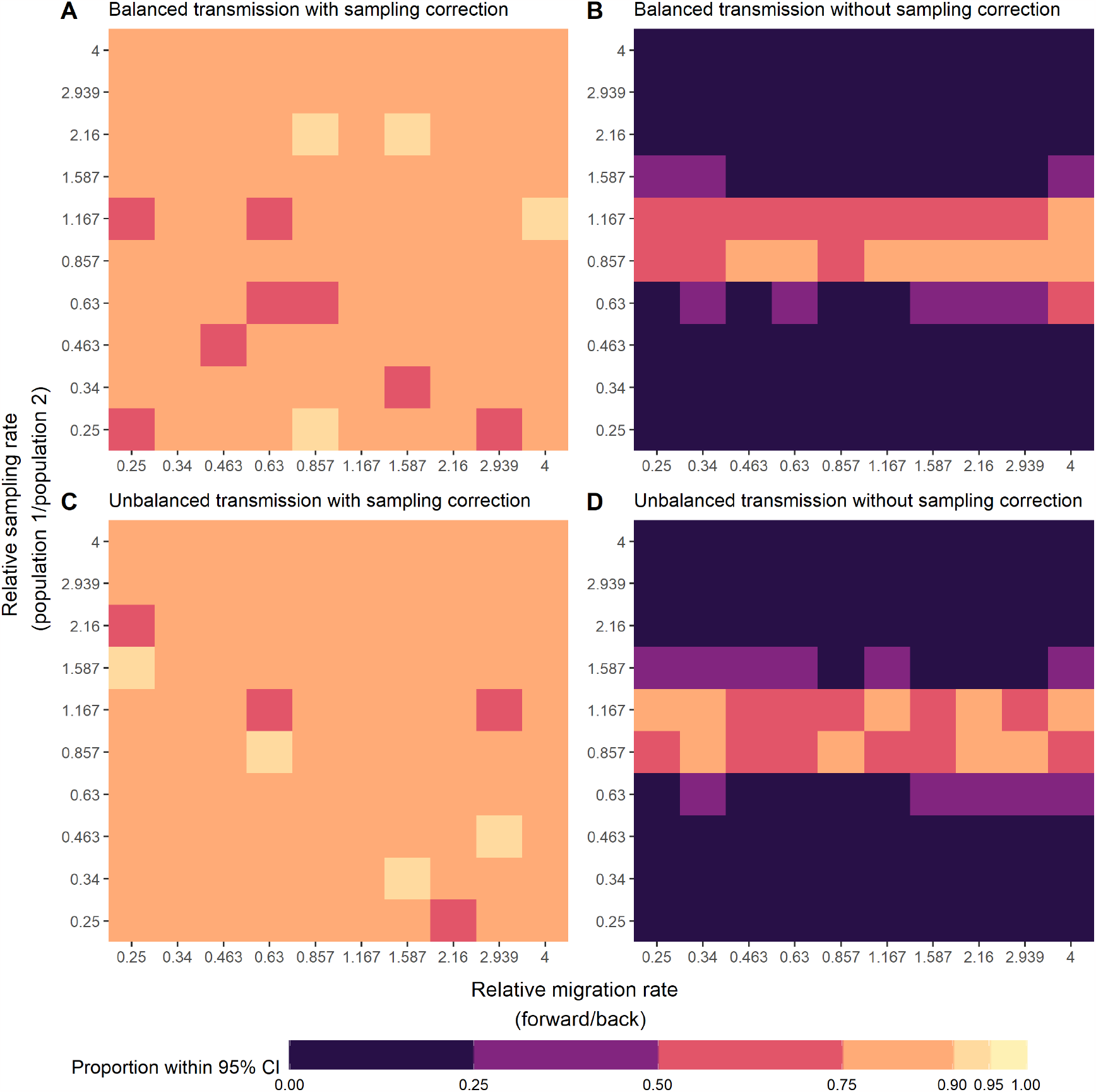
Proportion of model runs under the low sampling rate scenarios where the 95% confidence interval of the estimated relative migration rate includes the true relative migration rate from the simulation. We investigate different relative rates of migration and sampling to illustrate the performance of the proposed correction over a range of scenarios. These plots show the prediction under the corrected model (left; plots A and C) and the uncorrected all-rates different Mk model (Right; plots B and D). The top row (plots A and B) shows the performance of the correction and standard model in the simple case, where the transmission rate is equal across the two populations, and the bottom row (plots C and D) the biased transmission case, where transmission is two times higher in population 2 than population 1. Where a confidence interval could not be estimated due to the likelihood surface being flat in at least one direction the tree was dropped from the calculation.

Figure 4 shows that there is significant improvement in the confidence interval prediction with the correction. Where transmission rates are equal in the 2 populations, the true relative migration rate is included in the 95% CI of the predicted relative migration rate in the majority of simulations for every scenario with the correction. The minimum proportion of 95% CI predictions including the true value for any migration and sampling scenario is 0.68 compared to 0.00 with the uncorrected model. In the unequal transmission scenario, where transmission is 2 times higher in population 2 than in population 1, the correction continues to perform well with a minimum proportion of 0.73 of the reconstructed 95% confidence intervals including the true relative migration rate. Without the correction, this minimum value is again 0.00.

Under the transmission difference scenario, we see examples where very few tips are sampled from one of the two locations, as illustrated in Fig. 5. As migration rate reconstruction with DTA relies on the known location of sampled tips, it is much more difficult to infer a migration rate where one location is heavily under-represented in the phylogeny, but the correction appears to perform well at reconstructing the relative migration rate even in this scenario. In this scenario, however, a greater number of phylogenies are dropped due to inability to reconstruct 95% confidence intervals with both the corrected and uncorrected methods.

**Figure 5:**
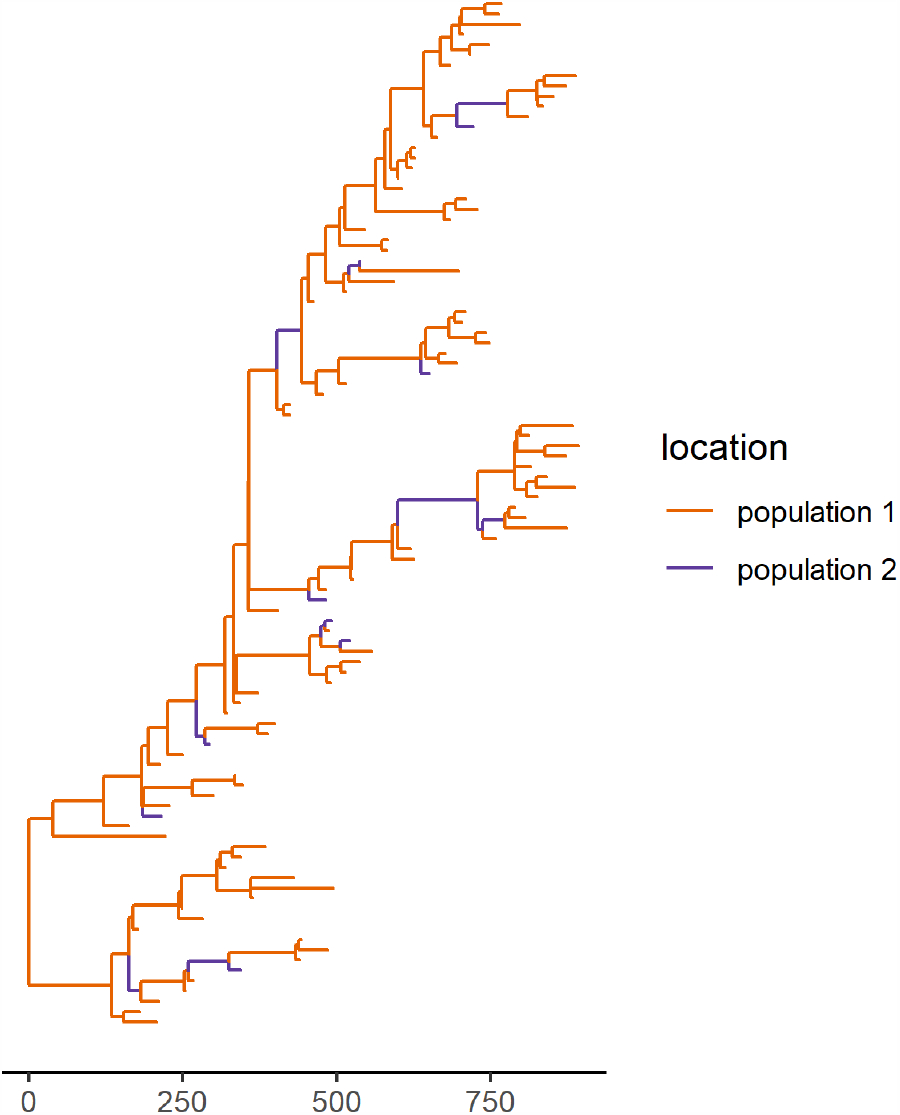
Example phylogeny for the scenario where migration from population 1 to 2 is 4 times lower than the reverse rate and sampling is 2.94 times lower (x= 0.34, y =0.25 on Fig. 4C) Branches are coloured according to their simulated location. This phylogeny is presented to highlight the lack of samples from population 2, making migration rate reconstruction difficult. The x-axis shows time in days from the root.

Figure 6 shows the MSLE for each modelled sampling and migration rate scenario. As with the confidence intervals, we see an overall improvement in prediction with the corrected method when sampling rates differ between locations. Under the equal transmission rate scenario, we see a highest MSLE of 0.33 with the correction and 2.56 without the correction (Fig. 6 A and B). In the transmission difference scenario, the correction performs similarly well, with a maximum MSLE of 0.32 with the correction and 2.43 without. We can also calculate the MSLE across the full range of true and predicted values in order to compare the corrected and uncorrected models. The MSLEs are the same for the transmission difference and low sampling rate scenarios with an overall MSLE of 0.07 with the correction and 0.86 without.

**Figure 6:**
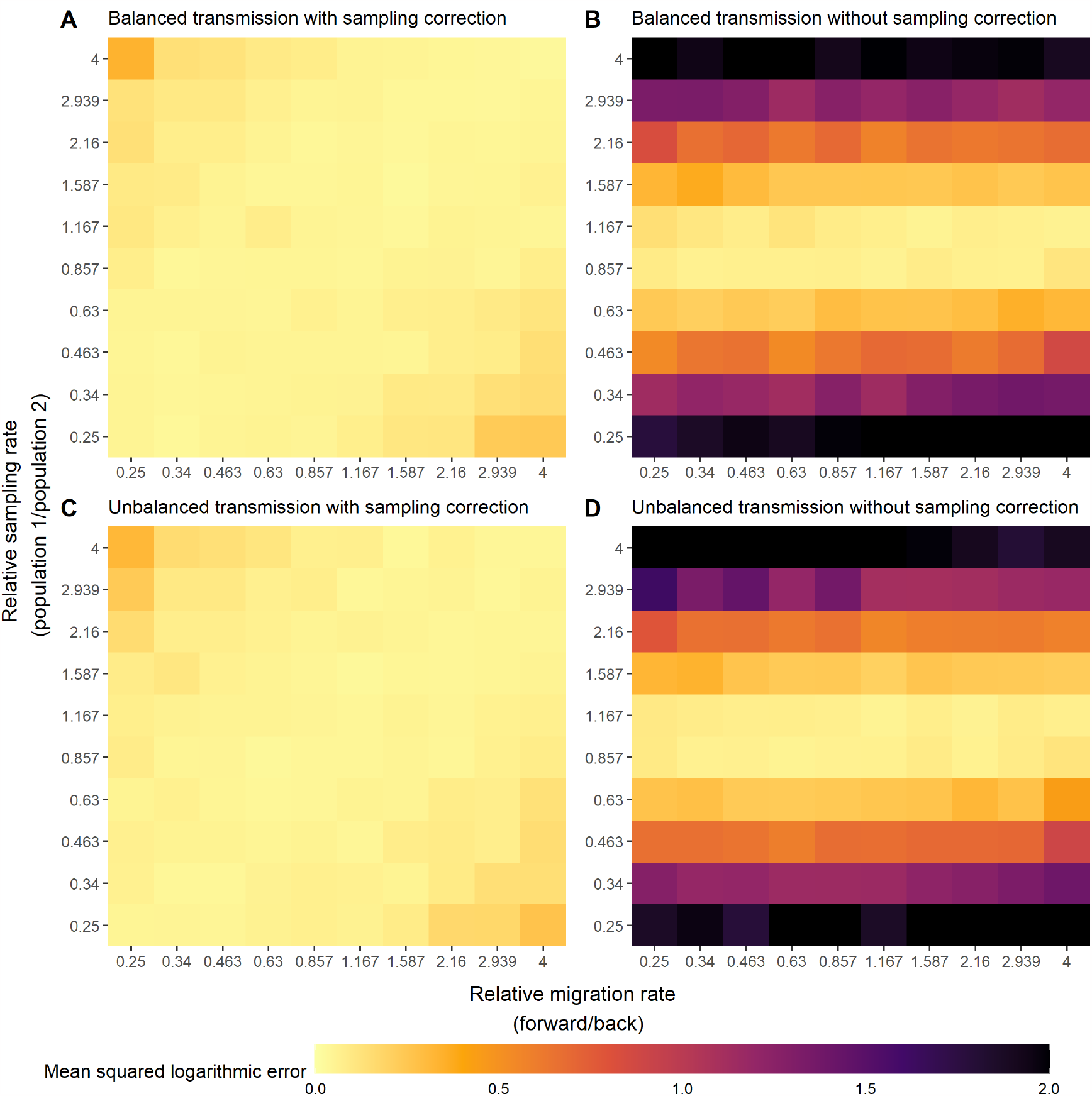
Difference between the true and estimated relative migration rates under the low sampling rate scenarios investigated. These plots show the mean squared logarithmic error (MSLE), a measure of the relative difference between the actual and predicted values. The left-hand panels (A and C) show the ability to reconstruct the true relative migration rate with the proposed correction and the right-hand panels (B and D) without. The top panels (A and B) show the performance where transmission rates are equal across the 2 populations and the bottom panels (C and D) where the transmission rate in population 2 is 2 times higher than in population 1. Values of the MSLE above 2 are shown in black on the above plot to allow better focus on the differences between lower values of the MSLE.

With the uncorrected method we see a clear horizontal band on the plots of both the proportion of 95% confidence intervals including the true value and MSLE for the predicted relative migration rates (Fig. 4 and 6). This suggests that the standard uncorrected method performs well at predicting the relative migration rate and the bounds on relative migration rate where the relative sampling rates are close to one. Further from the equal sampling case, where the differences in sampling rate are influencing the tip distribution of the phylogeny, the uncorrected method has much worse performance whereas the correction continues to perform well.

It is also evident that we see the worst performance with both the corrected and uncorrected methods where the phylogenies have very few tips from one location. This tends to occur most often where the rate of leaving a location is high and the rate of sampling in the same location is low or in the opposite case where the rate of leaving a location is low and the rate of sampling in that location is high (bottom right and top left of the plots in fig. 4 and 6).

#### 3.3.2 Medium sampling rate scenario

Under the medium sampling rate scenario, where sampling rates are on the same order of magnitude as migration rates, the correction performs similarly well (Fig 7) to in the low sampling scenarios above (Fig 6) with an overall MSLE of 0.08. We see an improvement with the uncorrected DTA in this scenario with an overall MSLE of 0.72. The corrected model performs significantly better than the uncorrected model when comparing overall MSLE (p *<* 0.001). 95% confidence intervals including the true relative migration rate and MSLEs under this scenario are shown in figure 7.

**Figure 7:**
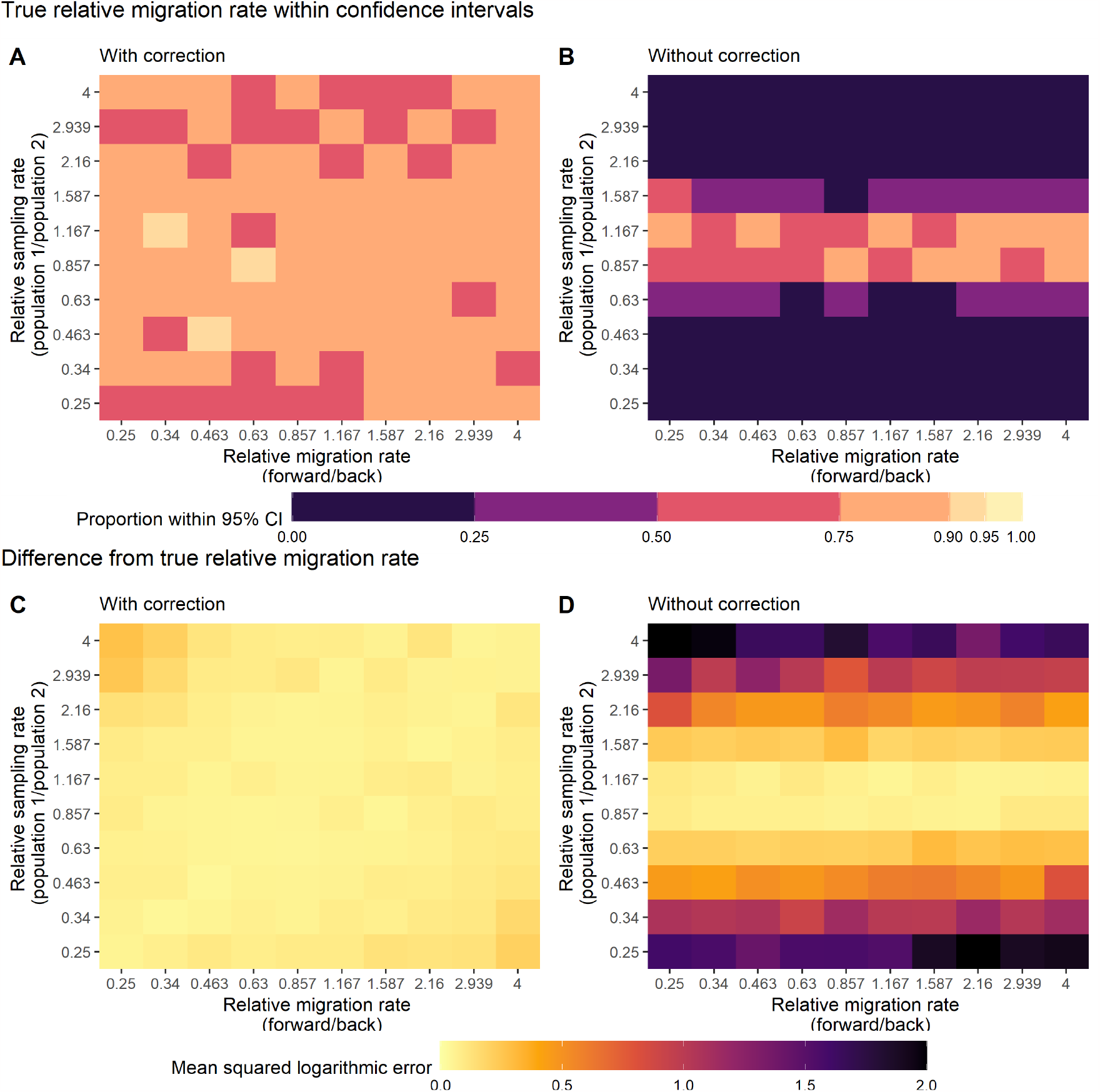
Predictive ability of the DTA model under the “medium sampling rate” scenario. A and B) Proportion of true relative migration rates within the 95% confidence intervals of the inferred relative migration rates under the corrected (A) and uncorrected (B) methods. C and D) Mean squared logarithmic error (MSLE), a measure of the relative difference between the actual and predicted values with the corrected (C) and uncorrected (D) methods. Values of the MSLE above 2 are shown in black on the above plot to allow better focus on the differences between lower values of the MSLE.

#### 3.3.3 High sampling rate scenario

With a high sampling rate, the sampled tips in the phylogeny are very close together in time and phylogenies tend to be rooted more recently than with lower sampling rates. In this case, the uncorrected DTA method is better able to reconstruct the relative migration rate than in the other sampling scenarios presented previously. The corrected (overall MSLE = 0.31) and uncorrected (overall MSLE = 0.29) methods perform equivalently (p=0.047). We posit that this is due to the much shallower phylogenies including fewer transmission events and less chance for alternative inference of migration events. In figure 8, with both the corrected and uncorrected methods, inference appears better close to the centre of the y-axis, where sampling rates are close to equal and sampling differences have less influence on the migration events. This suggests that sampling rate is still influencing the phylogenies produced. As both methods infer discrete location independently of tree topology, they are not able to correct for any topology differences introduced by biased sampling with high sampling rates.

**Figure 8:**
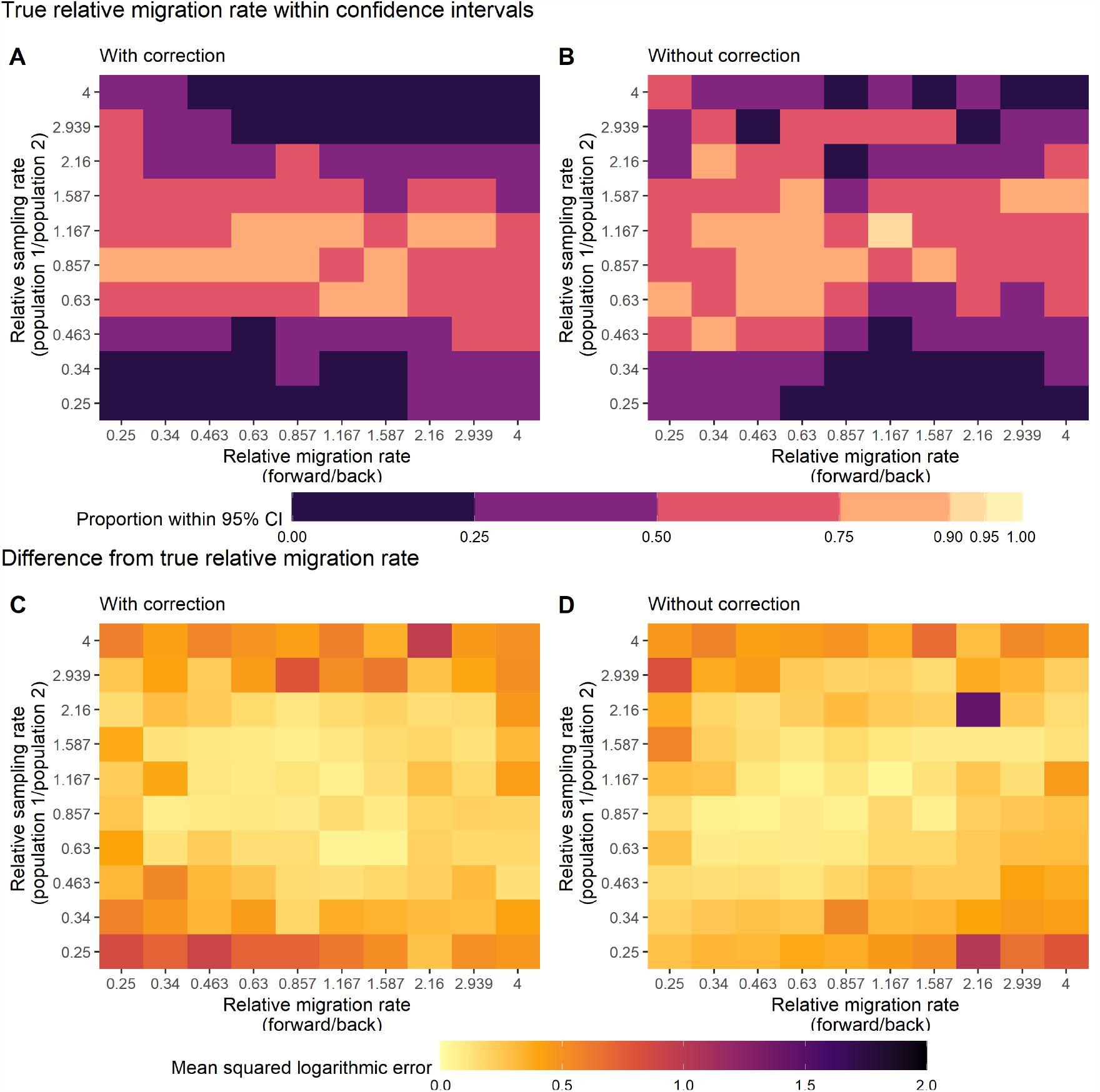
Predictive ability of the DTA model under the “high sampling rate” scenario. A and B) Proportion of true relative migration rates within the 95% confidence intervals of the inferred relative migration rates under the corrected (A) and uncorrected (B) methods. C and D) Mean squared logarithmic error (MSLE), a measure of the relative difference between the actual and predicted values with the corrected (C) and uncorrected (D) methods. Values of the MSLE above 2 are shown in black on the above plot to allow better focus on the differences between lower values of the MSLE.

## 4. Discussion

We have developed a correction for discrete trait phylogeographic analysis that improves estimates of migration rates between locations when sampling differs between them, thereby allowing more accurate inference of disease transmission across geographies. This correction can be simply implemented in standard phylogenetic software with minimal increase in computational time (mean of 0.19 seconds to reconstruct maximum likelihood migration rates and ancestral locations on a 100-tip, two-location phylogeny in the ace function from the R package ape compared to 0.12 seconds without the correction). The results presented in here show that the proposed correction can improve the inference of relative migration rates on phylogenies drawn from a simulation of an endemic disease where sampling rates are known, and vary by location. This is an important step as a major issue with the use of discrete trait analysis or mugration models is the assumption that sampling in a state is directly proportional to the underlying number of individuals in that state. In a two state scenario we show that the uncorrected Mk model, the base for standard discrete trait analysis, is only capable of reconstructing relative migration rates where sampling is approximately equal in each state as others have inferred previously [24, 47]. In the most biased low sampling rate scenario, the non-reversible DTA implemented fails to reconstruct a relative migration rate with a 95% confidence interval overlapping the true value in 100% of the 100 simulated phylogenies, we can reduce this to 42% with the proposed correction. This improves when larger sample sizes or larger numbers of phylogenies are considered.

As DTA continues to be widely used in phylogeography, it is important to consider and account for the limitations of the method, in the same way as approximations to the structured coalescent have been provided previously to account for the issues with speed of reconstruction and limitations on the number of states which can be feasibly considered [34]. The proposed correction is a first step, focusing on decoupling the sampling rate from the underlying population sizes. Several additional assumptions still exist with DTA which are at odds with traditional models of migration and can affect the tree structure and inferred parameters [24].

The proposed correction builds upon existing substitution models which account for the equilibrium frequency of bases with a similar method. The most widely used of these is the GTR model and using the weights in the GTR model to account for sampling rate differences when reconstructing geographic discrete traits has previously been suggested as a possibility [53]. The proposed amendment differs from the use of the GTR to model migration and sampling in two key ways: 1) The proposed amendment is based on a non-reversible transition rate matrix and 2) The amendment can be applied only at branches leading to sampled tips of the phylogeny. These differences are designed to better capture migration between discrete states on a short time scale, as is required when modelling geographic spread of viruses.

The correction can be easily implemented with code based on the ‘ace’ function from the ‘ape’ R package, including an eigen decomposition method which provides a significant fitting speed increase over performing matrix exponentiation on each fitting run. A Bayesian implementation of the proposed correction is presented on github (*https* : //*github*.*com*/*JorgensenD*/*PhD*_*t*_*hesis*) where both the tip-only and whole tree implementations are incorporated in a Markov Jump discrete trait analysis method. In future maximum likelihood use cases, selection of a 95% confidence interval estimation technique which is more robust to ridged likelihood surfaces or a transformation to produce a non-ridged surface would be preferable.

It is important to consider the limitations of the correction and of DTA and phylogeography more widely when selecting a method for different disease transmission scenarios. The method shown here is applied in an endemic scenario, where migration rate parameters are more simple to estimate as other parameters are steady over time. In an outbreak scenario, sampling and transmission rates can change dramatically over a short time period and previous studies have shown it can be hard to reconstruct migration history under these conditions [33]. This has been shown to also be the case with more complex structured coalescent models which have assumptions that can be inappropriate with complex population structures and emerging viruses and have been shown to perform more poorly than uncorrected DTA even without sampling rate differences [33, 54, 55]. The Mk model implemented here is applied on an empirical fixed set of simulated phylogenies. This highlights another issue with DTA, namely that the location data does not directly influence tree reconstruction and is, instead, inferred over a separately estimated phylogeny. As we consider sampling to have a significant effect on the migration rates estimated between locations, this bias is likely to also influence the depth and number of internal nodes estimated in the phylogeny itself. As a phylogeny only represents the relationships between the samples presented at the tips, any internal branching leading to unsampled lineages, for example, will not be included and unsampled locations will not be inferred at any ancestral nodes with DTA. Likewise, internal nodes along a branch are not included if both descendent lineages are not sampled. This can only be accounted for with a more complex, integrated method such as the structured coalescent or birth-death process. Sampling can also be geographically or phenotypically limited within each discrete region and not representative of the wider population, for example where environmental samples are only collected in a central city or from the outflows of a single hospital. This information can be lost when aggregating up to larger discrete regions and requires careful assessment of the discrete regions used or the treatment of location as a continuous rather than a discrete trait.

Continuous phylogeography offers the ability to infer an ancestral node outside of the sampled regions, an integrated inference of the tree and location, and produces dispersal velocities rather than migration rates between discrete regions which can be of interest when investigating the differential movement of virus clusters or the emergence of new viruses or serotypes [56]. Recent studies have indicated that continuous phylogeography can also be affected by differential sampling in space, although amendments to incorporate less certain spatial location data have been recently introduced [57, 39]. Current widely used continuous phylogeographic methods may be inappropriate for the use case of polio transmission, where distinct seasonal human migration patterns between rural and urban regions, distinct travel routes, religious movement events and nomadic populations within certain regions in the endemic Pakistan-Afghanistan region are at odds with the random Brownian motion viral movement assumption made. Different phylogeographic diffusion models developed by Barton, Etheridge and colleagues and outlined by Guindon et al. previously, have been implemented in recent continuous phylogeographic approaches [58, 57, 37]. These models are based on a generalised Wright-Fisher diffusion process known as the lambda-Fleming-Viot process, a model of extinction and recolonisation on a spatial continuum. The reverse of this model is the segregated lambda coalescent, allowing multiple lineages to coalesce at one time, rather than only two under the Kingman coalescent.[59, 60, 61].

Sub-sampling is a widely used alternative method to account for sampling biases in phylogenetic analysis. Here only a portion of the available data is included in analysis to produce a more representative dataset. Sub-sampling can be informed by a variety of demographic and epidemiological parameters such as the population size or number of reported cases in a region and recent work testing sampling schemes has been carried out on a set of Bayesian skyline methods [62]. Testing the same sub-sampling schemes on simulated structured population data and comparing the inferences under these sub-sampling schemes to the inferences with the proposed corrections would be of interest in the future, with improvement over sub-sampling vital to widespread adoption of the proposed correction.

Estimating the rate of genetic sampling for human viruses can often be challenging, especially where symptoms are not readily detected or are hard to differentiate from other infections. In reality, it is often not possible to know exact sampling rates and inferring them from collected data can be complex and introduce biases of its own. Where incidence and prevalence estimates can be made, via seroprevalence surveys or epidemiological modelling for example, the proportion of cases sequenced can be estimated and adjusted for by region. These processes introduce additional uncertainty not present under the modelled scenarios presented, further limiting the application of the proposed correction to real-world surveillance data.

The proposed correction provides a significant boost to the reliability of mugration model results where the sampling rate is known, particularly on a large number of phylogenies or where a large number of samples have been collected. As the sampling rates used are explicitly stated in the model, and there is little computational cost over standard maximum likelihood DTA, it would make sense to use the correction alongside the standard method to show the influence of potential sampling biases of different magnitudes on inferred migration rates. This comparison across a range of magnitudes of sampling bias between regions has been carried out in work to map the dispersal of HIV using a similar correction previously [63].

An example of a possible use case for the proposed correction is the phylogeographic analysis of polio sequences from acute flaccid paralysis data. As non-polio acute flaccid paralysis is monitored and reported globally, we expect the global distribution of cases to be approximately even according to population [64]. Differences from the expected non-polio paralysis rate by population size can be indicative of sampling differences and can be incorporated in the proposed correction when considering data sampled from polio paralysis cases.

As the results of DTA applied to migration are widely reported in literature, improving these methods without introducing a large amount of additional inference time or computational cost is important for phylogeography more widely and further research to address other flaws of the DTA method for migration is needed. Recent events also show a strong interest globally in phylogeography for outbreaks and epidemic disease and testing and modifying the proposed correction on simulated epidemic data would be a useful next step in development of more robust DTA models. Going forward it would also be useful to compare the inference under more complex structured coalescent and multitype birth-death methods to the proposed DTA correction, in addition to the comparison to standard DTA model presented here. A recent analysis by Seidel et al., and other previous studies, conclude that the birth-death and structured coalescent models perform similarly well in an endemic scenario with balanced sampling [65]. Comparison of these methods and the proposed correction under an endemic biased sampling scenario would be a useful comparison to make, as well as future extensions to incorporate more complex epidemic dynamics.

The proposed correction, although not able to address all of the issues with the use of DTA for migration rate inference, provides a significant boost to the precision of inferred migration rates in our simulated scenarios. As infectious disease genetic data collection expands, it is increasingly important to outline the limitations of specific methods and to provide updates and corrections to improve their inference ability.

### 4.1 Correction performance

As stated in the results section, the corrections with normalised sampling rates both performed better than the non-normalised sampling rate corrections when tested on simulated endemic data. This is likely due to the gradient descent (negative log likelihood minimisation) method used in this example. As normalising the sampling rates brings the base migration rates estimated (*q*_21_ in equation 8) onto a more similar scale to the relative migration rates for the reverse movements (*x* in equation 8), gradient descent updates are likely less dominated by the base migration rate. It would be interesting in the future to also rigorously test a true relative sampling rate with a reference population although this would require adaptations to existing model implementations rather than borrowing from the widely implemented equilibrium distribution of a DNA evolution model as we do here. The tip-only corrections perform better in these examples although further simulations with more clustered samples would be of interest to test the corrections on more heavily structured populations.

